# How Reliable are Prefrontal tDCS Effects – Zero–effects on the Keep–track Task?

**DOI:** 10.1101/736710

**Authors:** Dunja Paunović, Danka Purić, Jovana Bjekić, Saša Filipović

## Abstract

Recent neuroimaging studies showed that in addition to the dorsolateral prefrontal cortex (dlPFC), the posterior parietal cortex (PPC) plays a significant role in higher cognitive processes such as executive functions. In this study we aimed to explore the neural underpinnings of executive function of updating by exploring the effects of transcranial direct current stimulation (tDCS) over dlPFC and PCC. Nineteen healthy right–handed participants took part in a cross–over sham–controlled experiment. All participants underwent three tDCS conditions (active tDCS over the left dlPFC; active tDCS over the left PPC; and sham) in counterbalanced order. Following tDCS participants completed the keep–track task, with parallel forms being used in different test–sessions. As a control measure, we used a choice reaction time task. Results showed no significant effects of tDCS regardless of the localization of stimulation. Our results are in contrast with results of other studies exploring prefrontal tDCS effects on updating and do not allow for deriving conclusions about the role of the left PPC in the ability to update information in working memory.

## Introduction

Updating is defined as the ability to efficiently monitor incoming information and keep the relevant ones in mind by replacing older, no longer relevant with newer, more relevant information (Miyake et al., 2000; Morris & Jones, 1990; Purić, 2013). Not only by its definition, but also by its main operationalizations, such as n–back, keep–track, letter memory task. etc., this construct is similar to working memory (WM) (Miyake & Friedman, 2012; St Clair-Thompson & Gathercole, 2006; Wilhelm, Hildebrandt, & Oberauer, 2013).

### Neural basis of updating/WM

For many years, the functional activity of the dorsolateral prefrontal cortex (dlPFC) was considered to be the main hub for higher cognitive processes, especially executive functions (Ardila, 2018; Fuster, 2017; Levine, 2017). However, recent neuroimaging studies suggest that higher cognitive abilities depend on the activity of wider fronto–parietal functional network (Jung & Haier, 2007).

### Neuromodulation of Updating/WM

Neuromodulatory techniques enable exploration of causal relationships between the activity of certain brain regions and cognitive functions. One of the widely used is transcranial direct current stimulation (tDCS)–a noninvasive technique that modulates excitability of targeted brain areas by use of weak electrical current (Filmer, Dux, & Mattingley, 2014; Nitsche & Paulus, 2011).

The majority of tDCS studies of updating are mainly focused on anodal stimulation of the left dlPFC (Berryhill, Peterson, Jones, & Stephens, 2014; Dedoncker, Brunoni, Baeken, & Vanderhasselt, 2016). Even though most studies report facilitating effects of single–session tDCS over dlPFC (Dedoncker, et al., 2016), a certain number of studies show zero–effect (Andrews, Hoy, Enticott, Daskalakis, & Fitzgerald, 2011; M. Bogdanov & Schwabe, 2016; Hill, Rogasch, Fitzgerald, & Hoy, 2017, 2018; Mylius et al., 2012; Nilsson, Lebedev, & Lovden, 2015). However, regardless of their outcomes, all studies face some of methodological drawbacks and/or conceptual issues such as: treating a change in reaction times (RT) on the n–back task as a measure of updating, small sample sizes, cephalic positioning of reference electrode etc.

### Aims and Hypotheses

In this study we aimed to conceptually replicate the obtained prefrontal anodal tDCS effects on a different updating measure, i.e. the keep–track task which does not have the RT component, as well as to explore the effects of tDCS over PPC on updating so as to validate neuroimaging findings and derive possible causal relationship between the two. Finally, given the RT effects in previous studies, we wanted to assay the specificity of tDCS effects on updating. In line with these aims, we formulated the following hypotheses:

*H1*: *Stimulation over the left dlPFC will lead to increase in performance on the keep–track task*.

*H2*: *Stimulation over the left PPC will lead to increase in performance on the keep–track task*.

*H3: tDCS over dlPFC or PPC will not affect the performance on the choice reaction time task.*

## Method

### Design

This experiment was counterbalanced, randomized, sham–controlled, and employed cross–over factorial design. The within subject factor STIMULATION had two active (dlPFC and PPC) and one sham condition. We used accuracy on the keep–track and a choice RT task as the updating and the control measure, respectively.

### Participants

Sample size was determined *a priori* by power analysis for repeated measures with a three–level factor and parameters η^2^ = .15, α = .05, 1-β = .95. Based on the results we recruited 21 right–handed participants (age: *M* = 24.90, *SD* = 2.49; 11 female).

### tDCS

Three electrodes (5×5 cm) embedded in saline–soaked sponges were placed on participant’s scalp: two anodes (over F3 and P3 –International 10–20 EEG system), and the return over the right cheek (Figure 1).

**Figure 1:**
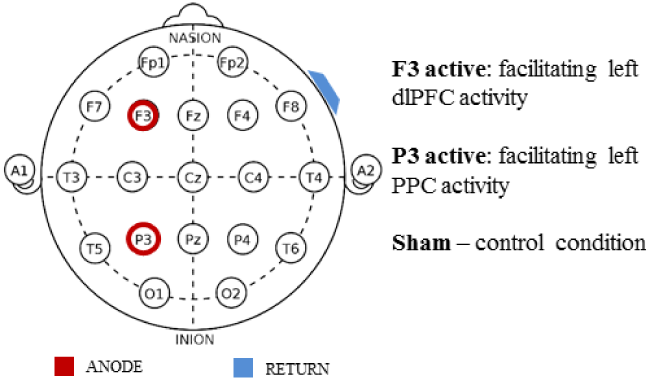
Electrode positioning

Active tDCS protocol included administering constant 1.5mA current for 20min with 30s fade– in/fade–out ramp using STMISOLA device (BIOPAC Systems, Inc, USA). In the first and the last minute of the sham protocol a 30s fade–in was immediately followed by the fade–out ramp.

### Tasks

In the keep–track task subjects monitor the list of successively presented 15 words. Each word in the list belongs to one of six semantic categories, but only representatives of the categories written in the bottom of the screen (*target categories*) should be payed attention to and recalled at the end of list presentation. There were total of nine lists. Number of target categories varied from three to five between lists. Accuracy was calculated as number of correctly recalled words (0-36).

In the choice RT task subjects gave consecutive responses to successive light up of four fields showed on the monitor by keypress. Each field had the corresponding key. Accuracy is measured as the average of RTs (time between light up and the keypress) for correct responses.

Parallel forms of both tasks were counterbalanced across sessions. The keep–track was administered right after and choice RT during the tDCS (Figure 2).

**Figure 2:**
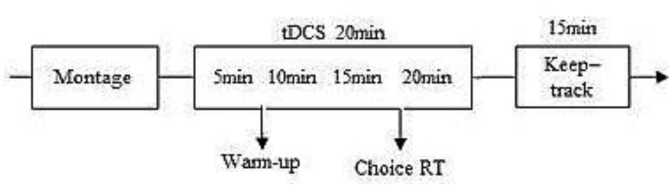
Session timeline

## Results

Statistical analyses were performed on the sample of 19 participants (two outliers), average age: *M* = 24.89 (*SD* = 2.60), 10 female. A post hoc power analysis with registered mean correlation between updating measures (*r =* .549) resulted in observed power 1-β = .987 adequate for performing planned analysis.

### tDCS effects on Updating

A repeated measures ANOVA with factor STIMULATION (F3/P3/sham) showed that scores on the keep–track task do not depend of presence or localization of stimulation (*F*_(2,36)_ = .499, *p* = .611) (Table 1).

**Table 1:** Descriptive statistics

| | | $M$ | $SD$ |
| --- | --- | --- | --- |
| Keep-track | F3 | 28.00 | 2.31 |
|  | P3 | 27.26 | 4.12 |
|  | Sham | 27.24 | 2.39 |
| Choice RT | F3 | 395.26 | 59.63 |
|  | P3 | 385.28 | 34.73 |
|  | Sham | 376.34 | 36.34 |

To exclude potential confounds such as TRAINING, GENDER and TASK FORMS, we performed additional analyses. A repeated measures ANOVA with factor TRAINING (first/ second/third session) showed marginal differences (*F*_(2,36)_ = 3.081, *p* = .058, η^2^ = .146) in participant’s scores between the first and the third session (*Post hoc* with Bonferroni correction: *p =*.055), while analyses for other two factors yielded insignificant results (mixed ANOVA: *F*_GENDER(1,17)_ = .035, *p* = .854; *F*_STIMULATION(2,36)_ = .462, *p* = .643; *F*_GENDERxSTIM(2,34)_ = .112, *p* = .894; repeated ANOVA: *F*_TASK_ FORMS(2,36) = 1.557, *p* = .225).

Effects of tDCS remained insignificant even when factor TRAINING was kept constant, as shown by a repeated measures ANOVA with factor STIMULATION on keep–track scores centered by session number (*F*_(2,36)_ = .921, *p* = .407).

### tDCS effects on control measure

A repeated measures ANOVA with factor STIMULATION showed that subject’s scores on the choice RT task are not modulated by tDCS (*F*_(1.281,23.059)_ = 1.519, *p* = .233) (Table 1).

## Discussion

This study shows that updating processes measured by the keep–track task are not modulated by tDCS regardless of its localization. These outcomes are in contrast with findings of positive prefrontal tDCS effects. However, most of these studies were based on the n–back paradigm (Berryhill & Jones, 2012; Fregni et al., 2005; Keeser et al., 2011) which leaves room for possibility that differences in outcomes originate from differences in task specific processes. On the other hand, our findings support those with zero effects (Andrews et al., 2011; Mario Bogdanov & Schwabe, 2016; Mylius et al., 2012; Nilsson et al., 2015) but all of these studies differ significantly in methodology and have setbacks that may call their validity into question. Furthermore, zero–effects on PPC do not allow assuming causality between PPC activity and processes engaged by the keep– track task.

Although our third hypothesis was confirmed, the pattern of results does not allow for conclusions about the specificity of tDCS effects but rather about the absence of random implausible effects.

Even though our results may call the reliability of prefrontal tDCS effects into question and do not allow for deriving conclusions about the role of the left PPC in the ability to update information they stress the importance of conducting more methodologically consistent research in order to validate either positive or zero–effects.

